# Promoting Human Intestinal Organoid Formation and Stimulation Using Piezoelectric Nanofiber Matrices

**DOI:** 10.1101/2024.06.12.598673

**Authors:** Holly M. Poling, Akaljot Singh, Maksym Krutko, Abid A. Reza, Kalpana Srivastava, James M. Wells, Michael A. Helmrath, Leyla Esfandiari

## Abstract

Human organoid model systems have changed the landscape of developmental biology and basic science. They serve as a great tool for human specific interrogation. In order to advance our organoid technology, we aimed to test the compatibility of a piezoelectric material with organoid generation, because it will create a new platform with the potential for sensing and actuating organoids in physiologically relevant ways. We differentiated human pluripotent stem cells into spheroids following the traditional human intestinal organoid (HIO) protocol atop a piezoelectric nanofiber scaffold. We observed that exposure to the biocompatible piezoelectric nanofibers promoted spheroid morphology three days sooner than with the conventional methodology. At day 28 of culture, HIOs grown on the scaffold appeared similar. Both groups were readily transplantable and developed well-organized laminated structures. Graft sizes between groups were similar. Upon characterizing the tissue further, we found no detrimental effects of the piezoelectric nanofibers on intestinal patterning or maturation. Furthermore, to test the practical feasibility of the material, HIOs were also matured on the nanofiber scaffolds and treated with ultrasound, which lead to increased cellular proliferation which is critical for organoid development and tissue maintenance. This study establishes a proof of concept for integrating piezoelectric materials as a customizable platform for on-demand electrical stimulation of cells using remote ultrasonic waveforms in regenerative medicine.

## Introduction

Human organoids are multicellular 3D structures that represent functional units of the larger organ and are derived from either adult or pluripotent stem cells^1–3^. Gastrointestinal (GI) organoids have been derived from definitive endoderm throughout the GI tract^4–11^. Organoids can be reconfigured to populate simple microfluidic devices or expanded with complex physiological properties approaching that of human tissue^5,12^. When a significant amount of mesenchyme co-develops with epithelium, these organoids can be readily transplanted to induce further maturation of the tissue with engraftment^7,8^. Because of these attributes, human organoids represent a great achievement in the way of modeling human organogenesis, development, physiology, and disease. However, various opportunities for enhancement persist, such as the capability to electrically stimulate them in real-time and promote their formation through a controllable and implantable platform.

Here, we focus on the human intestinal organoid (HIO) model system and incorporate a piezoelectric biomaterial during a critical window of the *in vitro* protocol as a feasibility study for the biocompatibility of organoids and engineered substrate constructs. Next, we employed the piezoelectric biomaterial to effectively stimulate maturing HIOs using on-demand ultrasound, thereby inducing a physiological response. The importance of electric fields has been well described during embryonic development, wound regeneration, and tissue homeostasis^13–16^. Exogenously applied electric fields have been shown to promote regeneration of tissues such as nerve, as well as other tissue regrowth^17–19^. These studies demonstrated that applying electric fields promoted enhanced nervous system regeneration, enhanced protein adsorption on surfaces^20^, bone regeneration^21,22^, angiogenesis^23,24^ and even phenotypic cellular changes^25,26^. Endogenous electric fields have also been an interesting topic within developmental biology for nearly a century^27–31^. Specific developmental events have been directly correlated to the generation of electric currents in embryonic animal model systems^32–37^. Of particular interest, substantial ionic currents exit the intestinal portal during the tail gut reduction period in chick embryos, where caudally extensive cell death occurs. Furthermore, disruptions in the intraembryonic voltage gradient was shown to cause developmental abnormalities with a 92% penetrance, demonstrating its critical importance^38^.

Piezoelectric materials provide a promising avenue for sensing and actuating, where a mechanical deformation of the material results in the endogenous polarization of ions or charges, inducing an electric potential^39–41^. In leveraging these unique material properties, we can facilitate organoid growth and maturation. Based on our understanding of electric fields in regeneration and development^14,15,17,42^ and their beneficial influence, we believe that piezoelectric material incorporation into HIOs can provide a platform to generate electric fields during organoid morphogenesis “on demand” through use of compatible stimulation, like ultrasound or shock waves instead of an invasive power source. Polyvinylidene fluoride-trifluoroethylene (PVDF-TrFE)^43^ is a synthetic polymer piezoelectric material that is both flexible and generally biocompatible. When electrospun, it can be manufactured into large pliable sheets with tunable primary fiber alignment and an overall thickness on the order of 10s of microns, making it an ideal, small, nanofiber based material to incorporate into *in vitro* organoid development. It can also be functionalized and modified to include a variety of biological molecules targeted at specific applications such as decellularized ECM^44,45^. While cells have been successfully cultured on PVDF-TrFE, to our knowledge, more complex biological systems have never been grown on these fibers^44,46–48^. Thus, in this study, we sought to determine whether organoid differentiation could be conducted on PVDF-TrFE membranes and if it could act in the stimulation of organoids under applied ultrasound.

## Materials and Methods

### Fabrication Procedure of PVDF-TrFE Scaffolds

PVDF-TrFE nanofiber scaffolds were electrospun as previously described^46^. Briefly, an electrospinning solution consisting of 20 % PVDF-TrFE (70/30) (PolyK Technologies) and solvent of N,N-dimethylformamide (DMF) and acetone (6/4 v/v) was made. The syringe pump’s flow rate was set to 1 mL per hour and a 20 G needle used. The needle tip was positioned 10 cm away from the drum collector which rotated at 2000 RPM throughout the process. This acted to produce fibers with a general linear alignment over a conductive polymer liner (McMaster-Carr). A voltage of 18 kV was applied between the needle tip and the collector. Three hour cycles were run producing scaffolds with an approximate thickness of 27 µm.

### Scanning Electron Microscopy

PVDF-TrFE scaffolds were stub mounted and sputter coated 12 nm thick with 60/40 gold palladium using an EM ACE600 (Leica). An SU8010 transmission electron microscope (Hitachi) was used to image samples.

### Animals

Male immunodeficient nonobese diabetic (NOD) severe combined immunodeficiency (SCID) interleukin-2Rγnull (NSG) mice aged between eight and ten weeks were used in all transplantation experiments. Mice were housed in the pathogen-free animal vivarium of Cincinnati Children’s Hospital Medical Center (CCHMC). Handling was performed humanely in accordance with the NIH Guide for the Care and Use of Laboratory Animals. NSG mice were fed antibiotic chow (275p.p.m. sulfamethoxazole and 1365p.p.m.trimethoprim; Test Diet). Both food and water were provided ad libitum pre- and post-operatively. All experiments were performed with the prior approval of the Institutional Animal Care and Use Committee of CCHMC (Building and Rebuilding the Human Gut, Protocol No. 2021-0060).

### Generation of HIOs

HIOs were generated and maintained as previously described with a substrate modification for one experimental group^4,49^. Briefly, H1 embryonic stem cells (WiCell Research Institute, Inc.) were grown in feeder-free conditions in Matrigel (BD Biosciences) coated plates and maintained with mTESR1 media (Stem Cell Technologies). For induction of definitive endoderm (DE), cells were plated at a density of 65,000 – 75,000 cells per well in 24-well plates. At this stage, cells were either plated in dishes alone, or in dishes containing a PVDF-TrFE mesh substrate. For both groups, cells were allowed to grow for two days before being treated with 100 ng/ml of Activin A for three days as previously described^4,7^. DE was then treated with hindgut induction medium (RPMI 1640, 100x NEAA, 2% dFCS,) for four days with 100 ng/ml FGF4 (R&D) and 3 µM Chiron 99021 (Tocris) to induce the formation of spheroids. Spheroids were then collected and plated in Growth Factor Reduced (GFR) Matrigel and maintained in intestinal growth medium (Advanced DMEM/F-12, N2 supplement, B27 supplement, 15 mM HEPES, 2 mM L-glutamine, penicillin-streptomycin) supplemented with 100 ng/ml EGF (R&D) to generate human intestinal organoids (HIOs). Thereafter, media was changed twice a week. HIOs were replated in fresh Matrigel after 14 days in culture. HIOs were utilized for surgical transplantation between days 28 and 32.

### Transplantation of Human Intestinal Organoids

A single HIO matured in vitro for 28-32 days was transplanted into the kidney capsule as previously described^7,50^. Mice were anesthetized with 2% inhaled isoflurane (Butler Schein) and the left flank was shaved and prepped in a sterile fashion with isopropyl alcohol and povidine-iodine. 1 cm posterior subcostal skin incisions were made, followed by retroperitoneal muscle incisions to expose the kidney. It was then removed from the cavity and a subcapsular pocket created. The individual HIO was implanted securely within the subcapsular pocket and secured at the distal end. Then, the kidney was returned to the peritoneal cavity and the incision was closed in a double layer. Upon closing, all mice were given a single injection of long acting Buprenex (0.05 mg/kg; Midwest Veterinary Supply) for pain management. Survival of mice was followed out to eight weeks at which time the mice were humanely euthanized.

### Gross measurement of HIOs

Transplanted HIOs were harvested and photographed alongside a metric ruler and analyzed using ImageJ (NIH). For every image, the engrafted HIO was measured widthwise and a 1 mm measurement on the ruler made in the same image was used as a conversion factor.

### Tissue processing, immunohistochemistry, and microscopy

Transplanted HIOs were harvested and fixed overnight in 4% paraformaldehyde at 4°C, processed and embedded in paraffin blocks. Sectioning was performed at a 5 μm thickness for staining. Slides were deparaffinized, rehydrated and antigen retrieval performed. Incubations with both primary and secondary antibodies took place overnight at 4°C. Antibodies and their respective dilutions are listed in Table 1. Slide based images were captured on a Nikon Eclipse Ti and analyzed using Nikon Elements Imaging Software (Nikon). For *in vitro* images, a Motic AE30 outfitted with a moticam 2300 was used (Motic Microscopes).

**Table 1.**
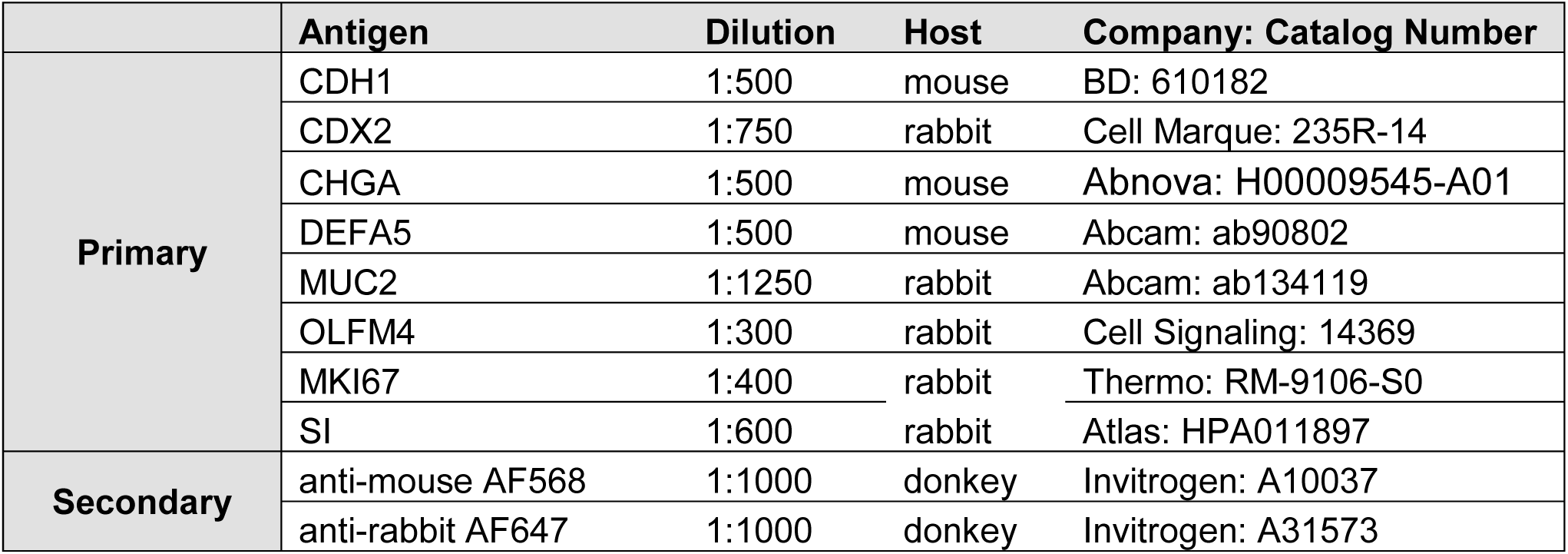
Antibodies and dilutions used for immunostaining.

### Morphometric Quantification

Morphometric analysis of the epithelial architecture was performed on hematoxylin and eosin stained tissue sections as previously described^10,51^. Crypt depth, villus height, and crypt fission were measured for a minimum of 20 well-oriented crypt-villus units per tissue sample and then averaged using Nikon NIS imaging software (Nikon).

### Bulk RNA Extractions and Sequencing

RNA was extracted from transplanted organoids using a RNeasy Plus Micro Kit (Qiagen) according to the manufacturer’s protocol. RNA samples were quantified and submitted to CCHMC’s DNA Sequencing and Genotyping Core for Next Generation Sequencing. All samples were assayed to have RNA integrity numbers greater than eight. After quality control, a cDNA library was created and sequenced using an IlluminaHiSeq2000 (Illumina) with 20 million paired-end reads per sample.

### Bulk RNA Sequencing Analysis

FastQC (v0.12.1) and MultiQC (v1.14) were used to assess the read quality of the fastq datasets. After aligning reads to human genome assembly hg38, they quantified using the quasi-mapper kallisto (v0.46.1) using default settings. Differential gene expression analysis was performed using R package DESeq2 (v1.38.3). Size factors were utilized to normalize the read count matrix, and a variance stabilizing transformation (VST) was applied to the normalized expression data. Sample correlation plot, volcano plot, dendrogram, and heatmap were generated using ggplot2 (v3.4.1) and heatmap (v1.0.12) packages. Gene ontology analysis was done using the ToppGene suite (toppgene.cchmc.org).

### Low-intensity Pulsed Ultrasound Treatments

A Sonitron GTS Sonoporation System (Nepagene) outfitted with a 12 mm diameter plane wave transducer was used to administer low-intensity pulsed ultrasound treatment. The transducer was positioned beneath the tissue culture plate directly below the growing organoids. A static holder was devised to reduce potential contact error arising from the manual manipulation of the transducer during treatment. Even contact was facilitated through a thin coating of aquasonic ultrasound transmission gel (Parker Laboratories, Inc.) between the transducer head and plate surface. Treatment parameters included an intensity of 0.3 W/cm^2^ of power constantly applied for 10 minutes.

### Proliferation Quantification

Tile scans of *in vitro* HIOs stained for Dapi, CDH1 and MKI67 were acquired on a Nikon Ti Eclipse (Nikon). Nikon Elements Software (Nikon) was used for smart thresholding and subsequent automated object counting. Each HIO’s nuclei counts expressing MKI67 were divided by the corresponding total nuclei counts and multiplied by 100 to yield the percent of MKI67+ cells in each organoid.

### Data Representation and Statistical Analysis

All graphs indicate the mean ± standard deviation. GraphPad Prism software (San Diego, CA, USA) was used to perform Student’s t-test and Wilcoxon-Mann-Whitney, based on distribution, for statistical analysis. A Fisher’s exact test was performed on binary data. For analysis including more than two experimental groups, an ANOVA followed by Tukey’s post-hoc analysis was performed. A p-value <0.05 was considered statistically significant.

## Results

### In Vitro Spheroid Generation

Thin flexible PVDF-TrFE scaffolds were fabricated through electrospinning for *in vitro* use (Supplementary Fig 1a). Scanning electron micrographs of the resultant scaffolds demonstrate the smooth nanofibers comprising the mesh produced by the manufacturing technique (Supplementary Fig 1b). In order to determine the compatibility of PVDF-TrFE with the generation of intestinal organoids, the scaffolds were implemented during the most critical window of HIO generation, hPSC differentiation and spheroid production (Supplementary Fig 2). HIOs were generated from multiple batches with and without PVDF-TrFE scaffolds until spheroid collection. The hPSCs cultured on PVDF-TrFE scaffolds exhibited phenotypic differences from those cultured without PVDF-TrFE (Fig 1a). Over the course of specification and differentiation, the cells on PVDF-TrFE appeared to grow as clusters and exhibit spherical conformations by day 3 , while those cultured without PVDF-TrFE scaffolds demonstrated this conformation starting at day 7. On day 9, spheroids were collected from both culture conditions, replated, and matured until day 28 in culture, at which time we observed the HIOs to appear similar between groups through bright field microscopy (Fig 1b).

**Figure 1.**
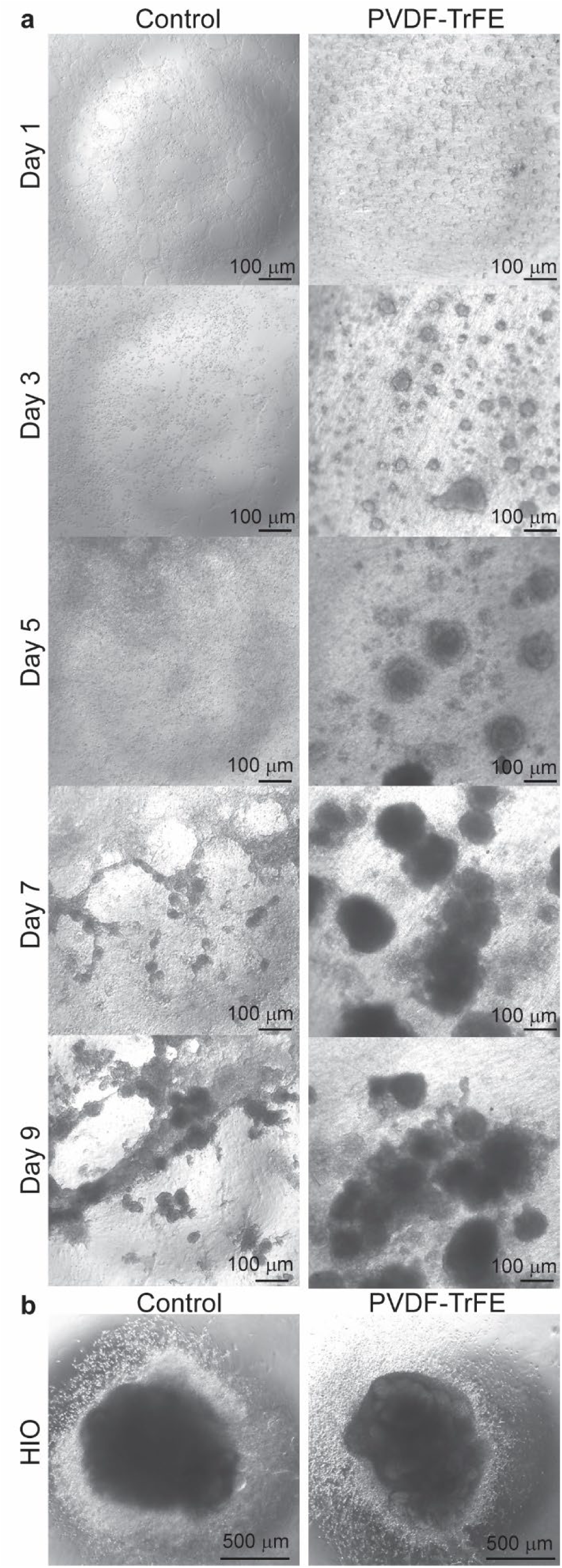
PVDF-TrFE scaffolds promote spheroid morphogenesis *in vitro* during HIO generation. **a**, Representative bright field images taken at days 1, 3, 5, 7, and 9 during the HIO generation protocol in Control and PVDF-TrFE groups. **b**, Representative bright field images of day 28 HIOs from control and PVDF-TrFE groups.

### Human Intestinal Organoid Engraftment

At the day 28 *in vitro* timepoint, HIOs from both groups were transplanted into the renal subcapsular space of immunocompromised (NSG) mice and allowed to engraft for eight weeks. We believed any significant differences would be highlighted best through the engraftment process and chose to focus on HIO characterization post-transplantation. After eight weeks of engraftment, transplanted HIOs (tHIOs), had a similar gross appearance between groups (Fig 2a). Upon quantifying the average tHIO graft width, we observed graft sizes to be similar between groups (Fig 2b). Histology of the tHIOs demonstrated the presence of key intestinal anatomical features in both groups (Fig 2c-d). When quantifying crypt fission, an indirect measure of stem cell activity in the epithelium of the samples, similar levels were found between groups (Fig 2e). When measuring both crypt depth and villus height across sample types, we also observed similarities between groups indicating that the epithelial surface area was well developed with and without PVDF-TrFE scaffolds during spheroid generation (Fig 2f-g).

**Figure 2.**
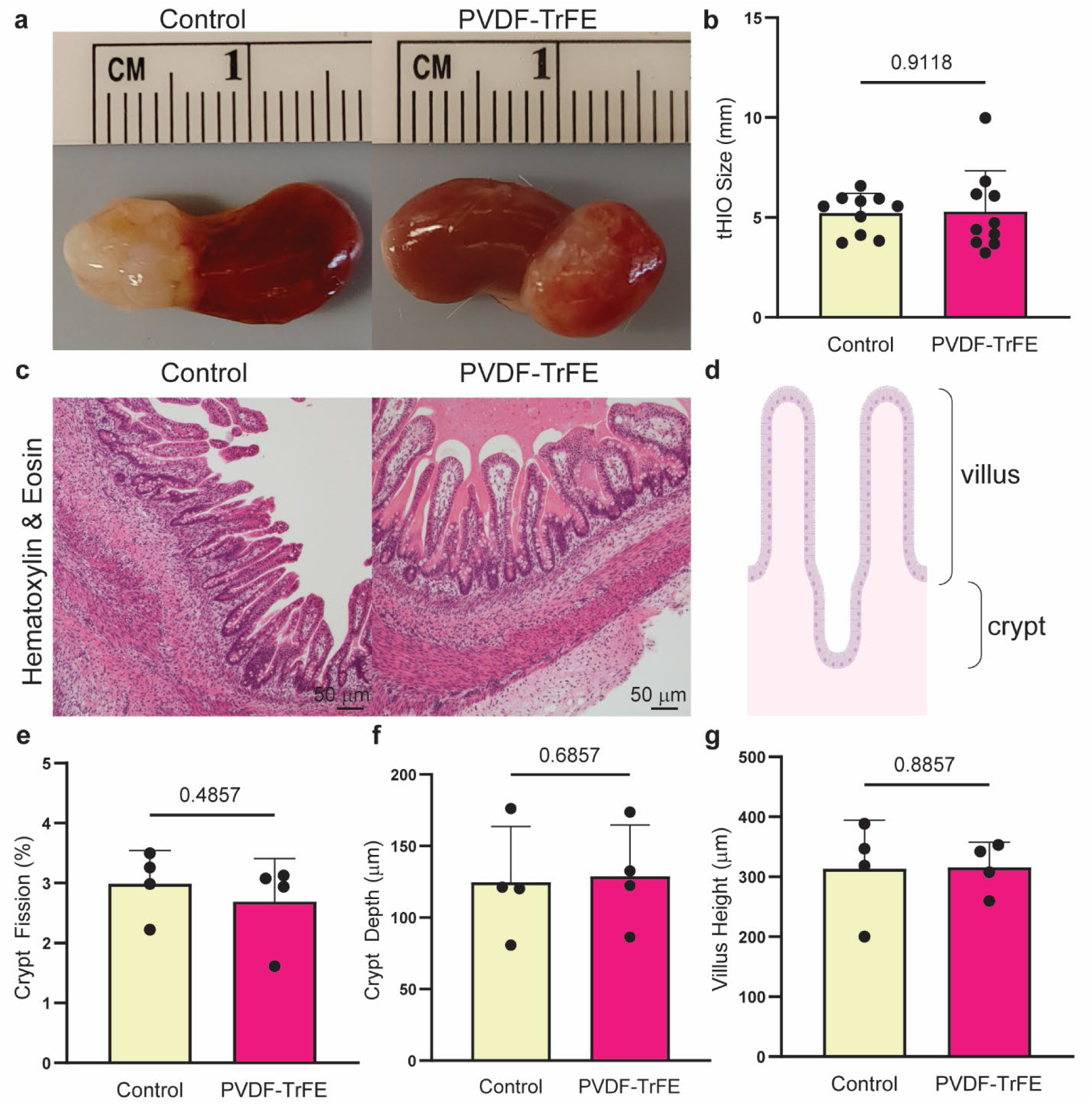
HIOs patterned on PVDF-TrFE readily transplant. **a**, Representative gross images of tHIOs engrafted under the kidney capsule of immunocompromised mice from control and PVDF-TrFE groups. **b**, Quantification of HIO graft size from images like those in a. tHIO size was similar between groups. n=6 per group. **c**, Representative hematoxylin and eosin stained sections of control tHIOs and those generated using a PVDF-TrFE scaffold. **d**, Intestinal architecture cartoon denoting Quantification of crypt depth between both groups was similar. **e**, Quantification of crypt fission between both groups was similar. **f**, Quantification of villus height between both groups was similar.

### Epithelial Characterization

Next, we interrogated epithelial cell type expression, which play important roles in digestion, nutrient absorption, and protection. Intestinal epithelial specificity was first confirmed by protein expression of marker Caudal Type Homeobox 2 (CDX2) in both groups (Fig 3a). Similarly, we demonstrated the presence of mucin producing goblet cells with Mucin 2 (MUC2) and a subset of hormone producing enteroendocrine cells with chromogranin a (CHGA) protein expression (Fig 3b). Paneth cell presence, cells important to host immunity, was also observed in both groups, as indicated by antimicrobial peptide alpha defensin 5 (DEFA5) expression in cells localized to crypt regions (Fig 3c). Crypts were further highlighted by staining for an antiapoptotic marker, olfactomedin 4 (OLFM4), and localized as expected in both groups (Fig 3d). Stereotypical proliferative zonation was observed through marker of proliferation KI67 (MKI67) expression suggesting epithelial renewal for maintenance and tissue homeostasis was achieved in both groups (Fig 3e). In order to gain functional insight, digestive enzyme presence was also validated. Sucrase isomaltase (SI), a dual-function enzyme for digestion of dietary starch, was strongly expressed apically on the epithelia in both groups (Fig 4a). Next, epithelial alkaline phosphatase activity, which plays an important role in gut mucosal defense, was examined by a precipitating reaction product and observed in both groups (Fig 4b).

**Figure 3.**
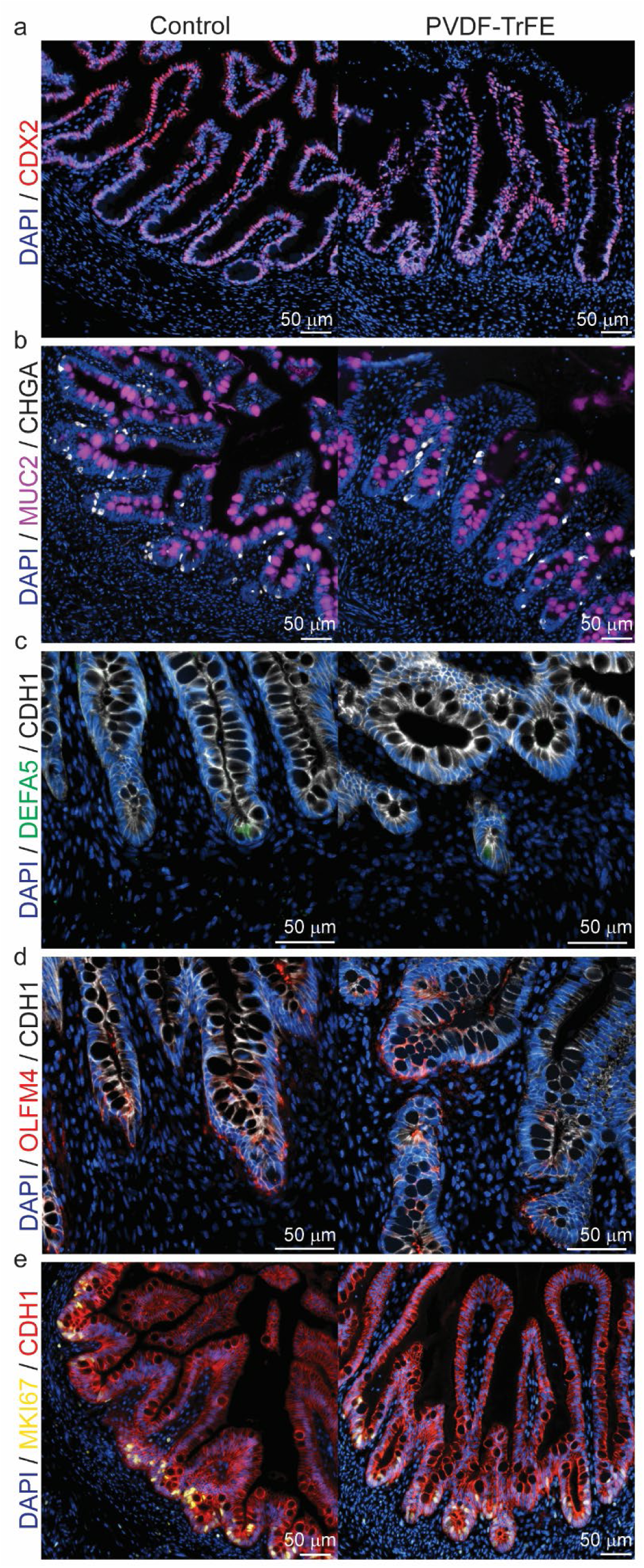
Epithelial cell type diversity is retained in HIOs generated on PVDF-TrFE. **a**, Representative immunostaining for intestinal epithelial cells (CDX2, red) and nuclei (DAPI, blue) in control and PVDF-TrFE groups. **b**, Representative immunostaining for goblet cells (MUC2, purple), enteroendocrine cells (CHGA, white) and nuclei (DAPI, blue) in control and PVDF-TrFE groups. **c**, Representative immunostaining for Paneth cells (DEFA5, purple), epithelium (CDH1, white) and nuclei (DAPI, blue) in control and PVDF-TrFE groups. **d**, Representative immunostaining for crypts (OLFM4, red), epithelium (CDH1, white) and nuclei (DAPI, blue) in control and PVDF-TrFE groups. **e**, Representative immunostaining for proliferation (MKI67, yellow), epithelium (CDH1, red) and nuclei (DAPI, blue) in control and PVDF-TrFE groups. n=3 per group.

**Figure 4.**
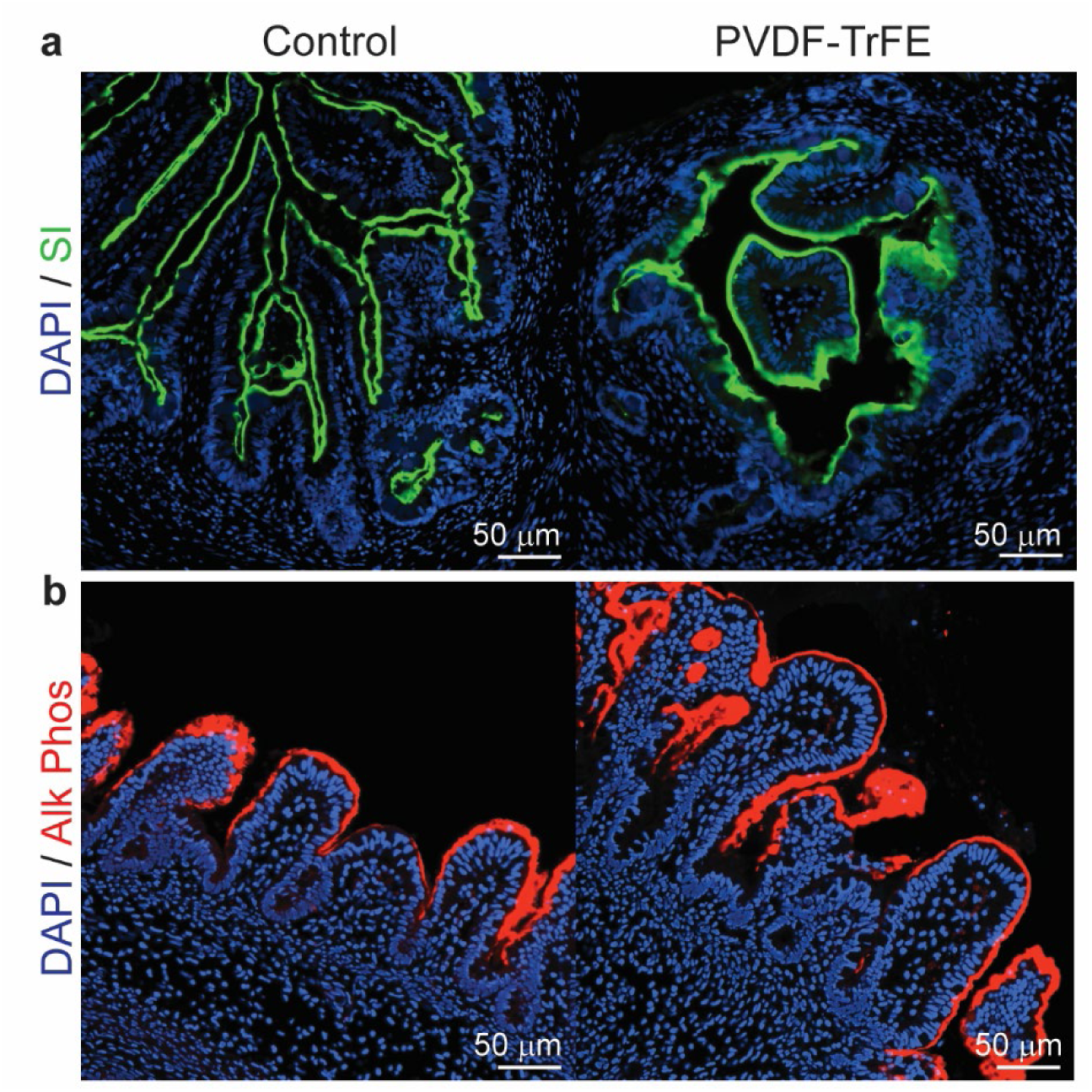
Digestive enzymes expression is retained in HIOs generated on PVDF-TrFE. **a**, Representative immunostaining for digestive enzyme sucrose isomaltase (SI, green) and nuclei (DAPI, blue) in control and PVDF-TrFE groups. **c**, Representative images demonstrating active alkaline phosphatase through precipitation (Alk Phos, red) and nuclei (DAPI, blue) in control and PVDF-TrFE groups.

### Transcriptomic Profiling

To investigate the organoids at the molecular level, bulk RNA sequencing was performed. Spearman correlation analysis of the samples revealed high coefficients for each pairwise comparison, all greater than 0.9, indicating a high degree of similarity between individual control and PVDF-TrFE scaffold differentiated organoid samples (Fig 5a). Furthermore, when constructing an unbiased dendrogram, samples from both experimental groups were mixed and the height of the branch points was low suggesting large similarities between samples (Supplementary Fig 3a). Of the 29,650 genes aligned and mapped, after filtering for read counts of five or greater in three samples, 17,524 genes were annotated. A volcano plot was used to visualize the differentially expressed genes between groups (Fig 5b). Only 37 protein coding genes were observed to be differentially expressed, which is further highlighted by a donut plot of the differentially expressed and non-differentially expressed genes between the two groups (Fig 5c). The differentially expressed genes are listed in Table 2, and did not strongly correlate to any specific biological processes. However, gene ontologies related to cellular components of the brush border were enriched, though genes from the input were low compared to those in the entire ontology annotation (Fig 5d). Furthermore, when examining a heatmap of curated genes related to important intestinal cell types, no striking differences were observed between groups (supplementary Fig 3b). Transcriptomically, the two groups demonstrate a substantial degree of uniformity with no remarkable differences. Together these data support the proper specification and development of intestinal tissues with the use of PVDF-TrFE as a scaffold during spheroid generation.

**Figure 5.**
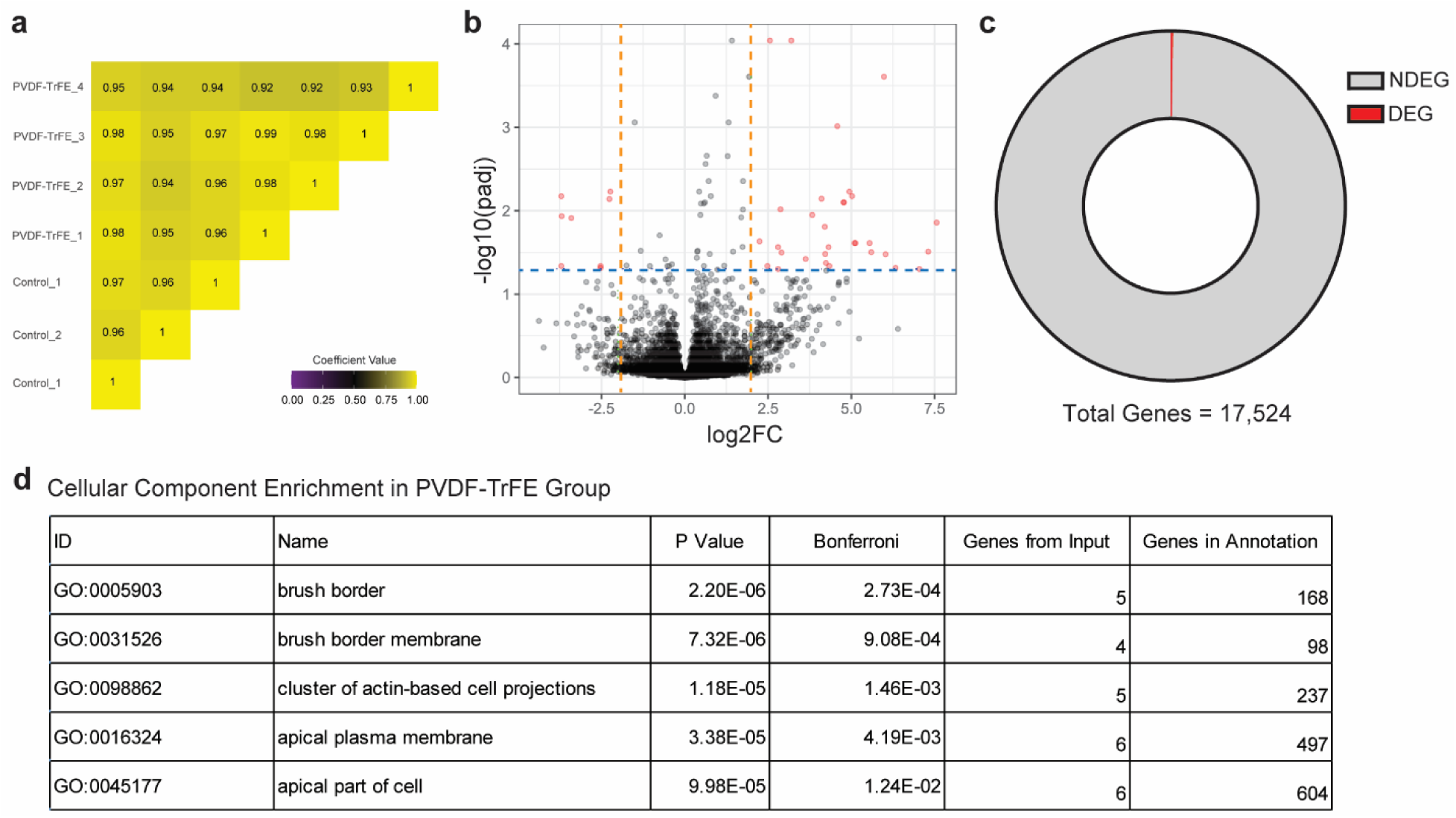
Transcriptomic profiles are similar in HIOs generated with and without PVDF-TrFE scaffolds. **a**, Spearman correlation matrix of bulk RNA sequencing data obtained from control HIOs and PVDF-TrFE HIOs after transplantation. **b**, Volcano plot of bulk RNA sequencing data obtained from control HIOs and PVDF-TrFE HIOs after transplantation with orange dotted lines indicating the fold change cutoff and blue dotted line indicating the significance threshold of p=0.05. **c**, Donut chart quantification of the differentially and non-differentially expressed genes as demonstrated in b. **d**, Uncurated table of the top five enriched cellular compartment gene ontologies in HIOs generated with PVDF-TrFE compared to HIOs generated without PVDF-TrFE.

**Table 2.** Differentially Expressed Genes in Transplanted HIOs.

| Gene | log2FoldChange | P Value | Adjusted P Value | Ensembl Gene ID |
| --- | --- | --- | --- | --- |
| GOLGA8O | 7.562 | 2.775E-05 | 1.389E-02 | ENSG00000206127 |
| ATP6V0A4 | 5.981 | 6.082E-08 | 2.474E-04 | ENSG00000105929 |
| SLC34A1 | 5.599 | 8.972E-05 | 3.144E-02 | ENSG00000131183 |
| CYP2A7 | 5.549 | 5.682E-05 | 2.441E-02 | ENSG00000198077 |
| CYP4A11 | 5.103 | 5.536E-05 | 2.441E-02 | ENSG00000187048 |
| MIOX | 4.940 | 5.099E-06 | 5.904E-03 | ENSG00000100253 |
| SLCO1A2 | 4.580 | 4.971E-07 | 9.678E-04 | ENSG00000084453 |
| G6PC1 | 4.343 | 1.812E-04 | 4.601E-02 | ENSG00000131482 |
| UGT1A6 | 4.320 | 6.709E-05 | 2.728E-02 | ENSG00000167165 |
| PDZK1IP1 | 4.219 | 9.996E-05 | 3.305E-02 | ENSG00000162366 |
| CYP4B1 | 4.201 | 3.202E-05 | 1.559E-02 | ENSG00000142973 |
| GPX3 | 4.105 | 8.596E-06 | 7.173E-03 | ENSG00000211445 |
| ACSM2A | 3.825 | 1.987E-05 | 1.123E-02 | ENSG00000183747 |
| SLC26A4 | 3.628 | 1.211E-04 | 3.790E-02 | ENSG00000091137 |
| KCNJ16 | 3.197 | 8.900E-09 | 9.106E-05 | ENSG00000153822 |
| SMYD1 | 2.906 | 9.244E-05 | 3.176E-02 | ENSG00000115593 |
| PAH | 2.874 | 1.540E-05 | 9.641E-03 | ENSG00000171759 |
| RSC1A1 | 2.802 | 2.082E-04 | 4.999E-02 | ENSG00000215695 |
| SLC27A2 | 2.802 | 6.851E-05 | 2.728E-02 | ENSG00000140284 |
| NRXN3 | 2.557 | 1.559E-08 | 9.106E-05 | ENSG00000021645 |
| KMO | 2.482 | 1.694E-04 | 4.601E-02 | ENSG00000117009 |
| P2RY14 | 2.248 | 5.045E-05 | 2.327E-02 | ENSG00000174944 |
| LRRC55 | 1.932 | 7.058E-08 | 2.474E-04 | ENSG00000183908 |
| ACMSD | 1.748 | 3.519E-06 | 4.404E-03 | ENSG00000153086 |
| DGKG | 1.746 | 1.607E-05 | 9.709E-03 | ENSG00000058866 |
| MGAM | 1.733 | 6.509E-05 | 2.716E-02 | ENSG00000282607 |
| GK | 1.694 | 2.264E-05 | 1.202E-02 | ENSG00000198814 |
| TNMD | -1.503 | 3.595E-07 | 8.733E-04 | ENSG00000000005 |
| SLFN12 | -1.746 | 1.552E-04 | 4.532E-02 | ENSG00000172123 |
| CHODL | -2.231 | 5.728E-06 | 5.904E-03 | ENSG00000154645 |
| CPA6 | -2.259 | 9.077E-06 | 7.230E-03 | ENSG00000165078 |
| STRA6 | -2.525 | 1.793E-04 | 4.601E-02 | ENSG00000288257 |
| OSR2 | -2.529 | 1.971E-04 | 4.865E-02 | ENSG00000164920 |
| SYT10 | -3.403 | 2.375E-05 | 1.224E-02 | ENSG00000110975 |
| ZIC1 | -3.693 | 2.120E-05 | 1.161E-02 | ENSG00000152977 |
| MYBPC1 | -3.704 | 1.775E-04 | 4.601E-02 | ENSG00000196091 |
| CXCL6 | -3.704 | 7.652E-06 | 6.704E-03 | ENSG00000124875 |

### Ultrasound Mediated Piezoelectric Material Activation for Electric Field Production

Once we were convinced that PVDF-TrFE was compatible with the *in vitro* HIO model system, we sought to test its practical application. To do this, we differentiated spheroids in the conventional manner and re-plated them atop PVDF-TrFE nanofiber scaffolds (Supplementary Fig 4a). Then, after continued culture, low-intensity pulsed ultrasound (LIPUS) treatments were administered every other day for the last week of culture. Treatments were 10 minutes in duration with 0.3 W/cm^2^ of power applied through a plane wave transducer (Supplementary Fig 4b). Previous reports demonstrated a positive impact of this treatment level on the proliferation of homogeneous cell line culture systems, so we chose it for our experiment^52–57^. To interrogate proliferation, MKI67 immunofluorescence was performed on four experimental groups of HIOs: control, without PVDF-TrFE as a substrate, with PVDF-TrFE as a substrate and then with and without LIPUS treatment (Fig 6a). When quantifying the percent of MKI67+ cells across all four groups, we observed several substantial differences (Fig 6b). In general, across all groups, it appeared that most MKI67+ cells were CDH1-, and therefore part of the developing mesenchymal compartment. We then confirmed that the LIPUS treatment regimen alone had a positive effect on HIO proliferation, and that PVDF-TrFE as a substrate positively impacted HIO proliferation. Furthermore, in the case of growing HIOs on PVDF-TrFE with LIPUS treatments, an additive proliferative effect was observed. This group demonstrated the highest percentage of MKI67+ cells suggesting that a piezoelectric material can serve both alone and in conjunction with LIPUS stimulation of organoids to produce proliferative effects.

**Figure 6.**
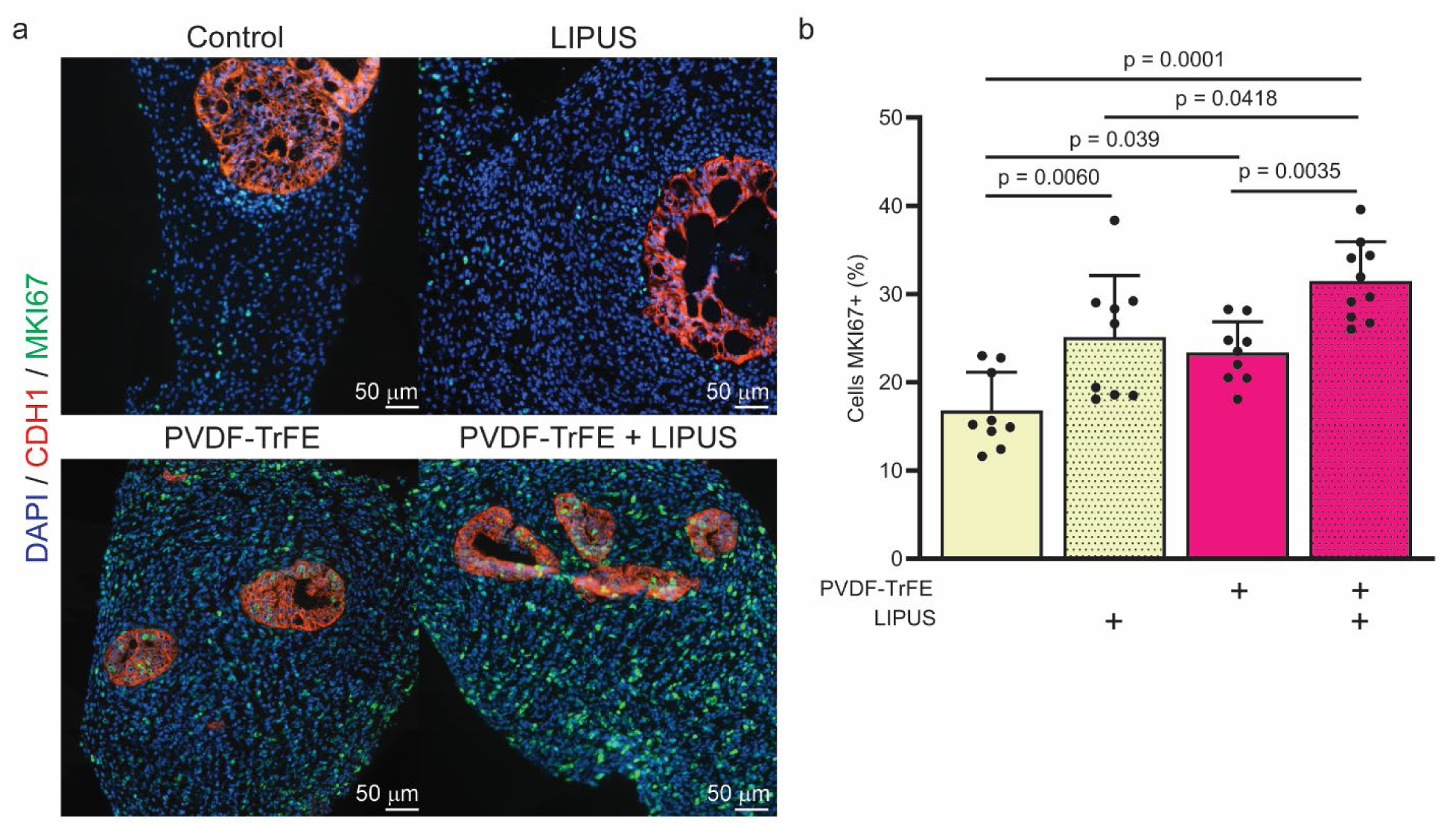
PVDF-TrFE activation with LIPUS increases HIO proliferation. **a**, Representative immunofluorescence staining for proliferation (MKI67, green), epithelium (CHD1, red), and nuclei (DAPI, blue) in HIOs cultured with and without PVDF-TrFE as a substrate and with and without LIPUS treatment. **b**, Quantification of proliferation (MKI67+ cells) from tile scans of staining in a.

## Discussion

HIOs can be successfully generated when using PVDF-TrFE as a differentiation substrate. Through the patterning process, spheroid formation was observed 4 days sooner with the use of PVDF-TrFE when compared to control. After 28 days in vitro, the HIOs from both groups were grossly similar. Eight weeks after transplantation, control HIOs and PVDF-TrFE HIOs grew to similar sizes. The resultant intestinal architecture and morphometric features was also similar between groups. Differentiated epithelial cell types were well represented in both groups and the characteristic intestinal epithelial proliferative zonation for self-renewal and homeostasis was observed in both groups. The presence of digestive enzymes was also observed to be similar with and without PVDF-TrFE. Transcriptomic profiles between both groups of resultant HIOs were also highly uniform. Taken together, these data suggest that proper cell fate decisions and overall intestinal patterning can readily proceed in the presence of PVDF-TrFE.

This study serves as a proof of principle for the incorporation of piezoelectric materials into organoids and opens the door to a new generation of HIOs. Because we did not observe any detrimental effects of PVDF-TrFE during the most sensitive stages of making an HIO, we conclude that it is truly biocompatible with developing complex cell structures. In our pilot study, we also demonstrated that using a piezoelectric material as a substrate can enhance organoid proliferation which can be augmented by LIPUS activation of the substrate material. Several recent studies have focused on the incorporation of nanoelectronics and brain organoids, primarily for sensing voltages, however incorporating both sensing and actuator capabilities into such a setting remains an opportunity^58–61^.This opportunity can be realized through the application of multipurpose piezoelectric materials. For modeling organs with inherent mechanical activity like those throughout the gastrointestinal tract, this is especially helpful as we may soon be able to simulate churning, segmentation, and peristalsis functions *in vitro* for increased organoid maturation. We may also begin to elucidate the collective behavior of multi-organoid and large-scale organoid structures to gain insight on the forces of organogenesis through measuring the electrical output of the piezoelectric fibers as a sensor. Furthermore, because these materials are biocompatible, the potential to incorporate them with transplantation also exists. In our continued line of study, we aim to accomplish *in vivo* organoid actuation. A diverse range of experimentation will be made possible through ‘piezorganoids’ and the marriage of engineering and developmental biology.

## Acknowledgements

Research reported in this publication was supported by National Institute of Diabetes and Digestive and Kidney Disorders (NIDDK) and National Institute of Allergy and Infectious Diseases (NIAID) of the National Institutes of Health under grant number U01DK103117 (MAH, JMW). The content is solely the responsibility of the authors and does not necessarily represent the official views of the National Institutes of Health. Additional support was provided by the National Science Foundation Division of Materials Research under grant number DMR-2104639, and the University of Cincinnati as a startup research funding (LE). Additional financial support was provided by a National Science Foundation Graduate Research Fellowship Program award (HMP). The authors thank Yara Izhiman for 3D printing support. Figure schematics and cartoons were created with Biorender.com.

## Author Contributions

H.M.P. and M.A.H. and L.E. conceived and designed the study; H.M.P., A.S, M.K. and K.S. performed experiments; H.M.P., A.S., and A.A.R., analyzed data; H.M.P., A.S., A.A.R., J.M.W., M.A.H., and L.E. interpreted experimental findings; H.M.P. prepared figures and drafted manuscript; all authors edited the manuscript and approved the final version.

## Data Availability

The authors declare that all data supporting the findings of this study are available within the paper and its supplementary information. Sequencing data is available at ArrayExpress accession E-MTAB-13578. Supplementary information is available for this paper. Correspondence and requests for materials should be addressed to Michael A. Helmrath or Leyla Esfandiari.

---

**Supplementary Figure 1.**
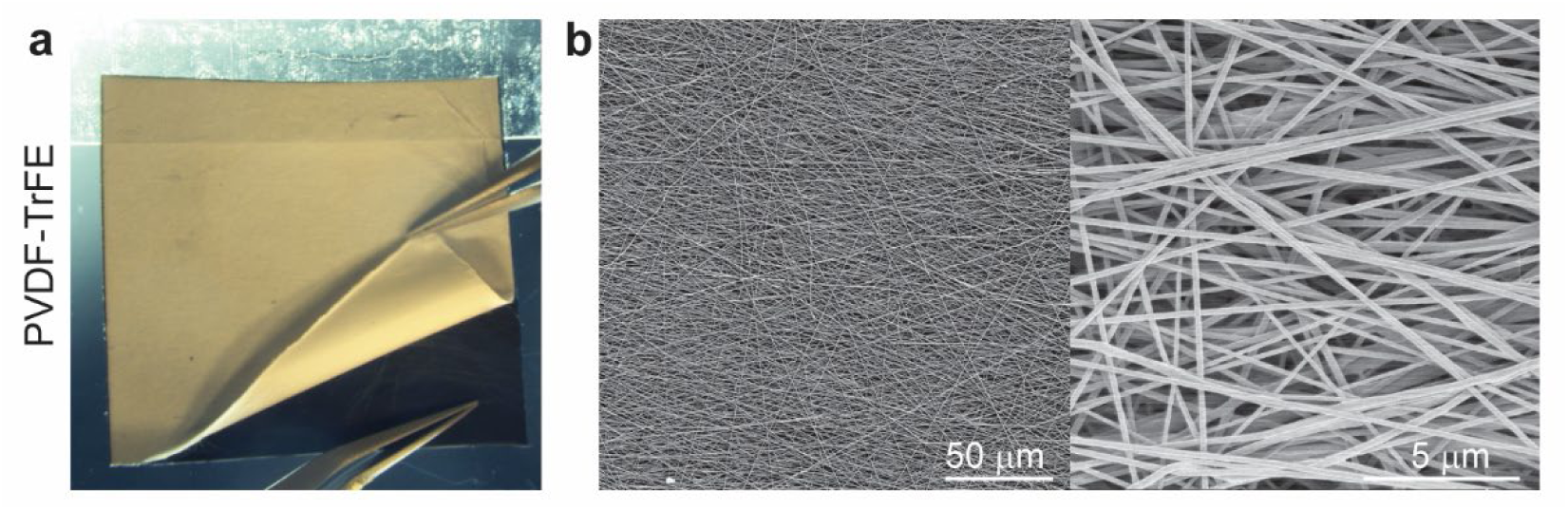
PVDF-TrFE material characterization. **a**, Photograph of 3 hour electrospun PVDF-TrFE being removed from its backing. Note the flexibility of the mesh. **b**, Scanning electron micrographs at low and high magnification demonstrating the smooth fibrous structure of the scaffolds.

**Supplementary Figure 2.**
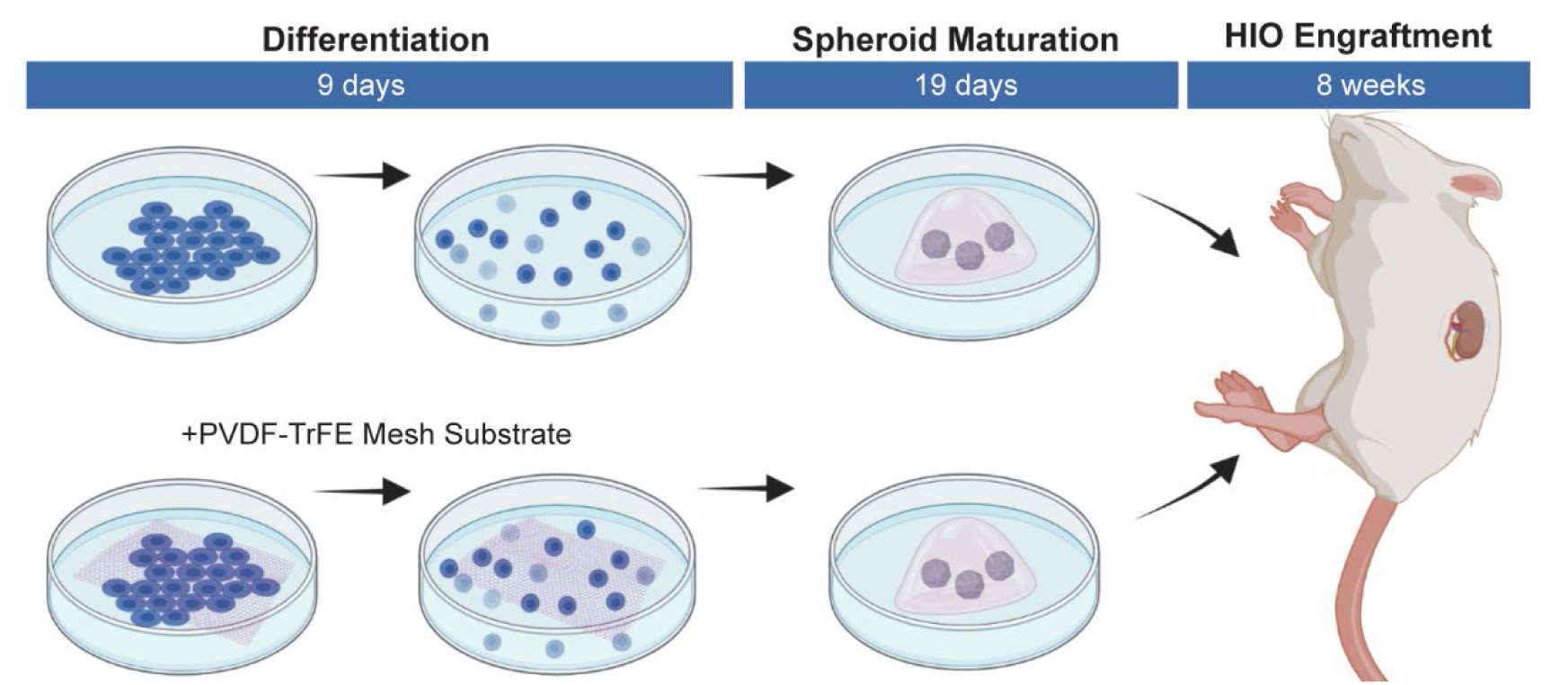
Experimental schematic for in vivo transplantation experiment. Human pluripotent stem cells were seeded into well plates following conventional methods or atop a PVDF-TrFE nanofiber scaffold for standard differentiation. After spheroids were generated and collected, they were plated normally and continued to mature for 28 days in culture prior to transplantation.

**Supplementary Figure 3.**
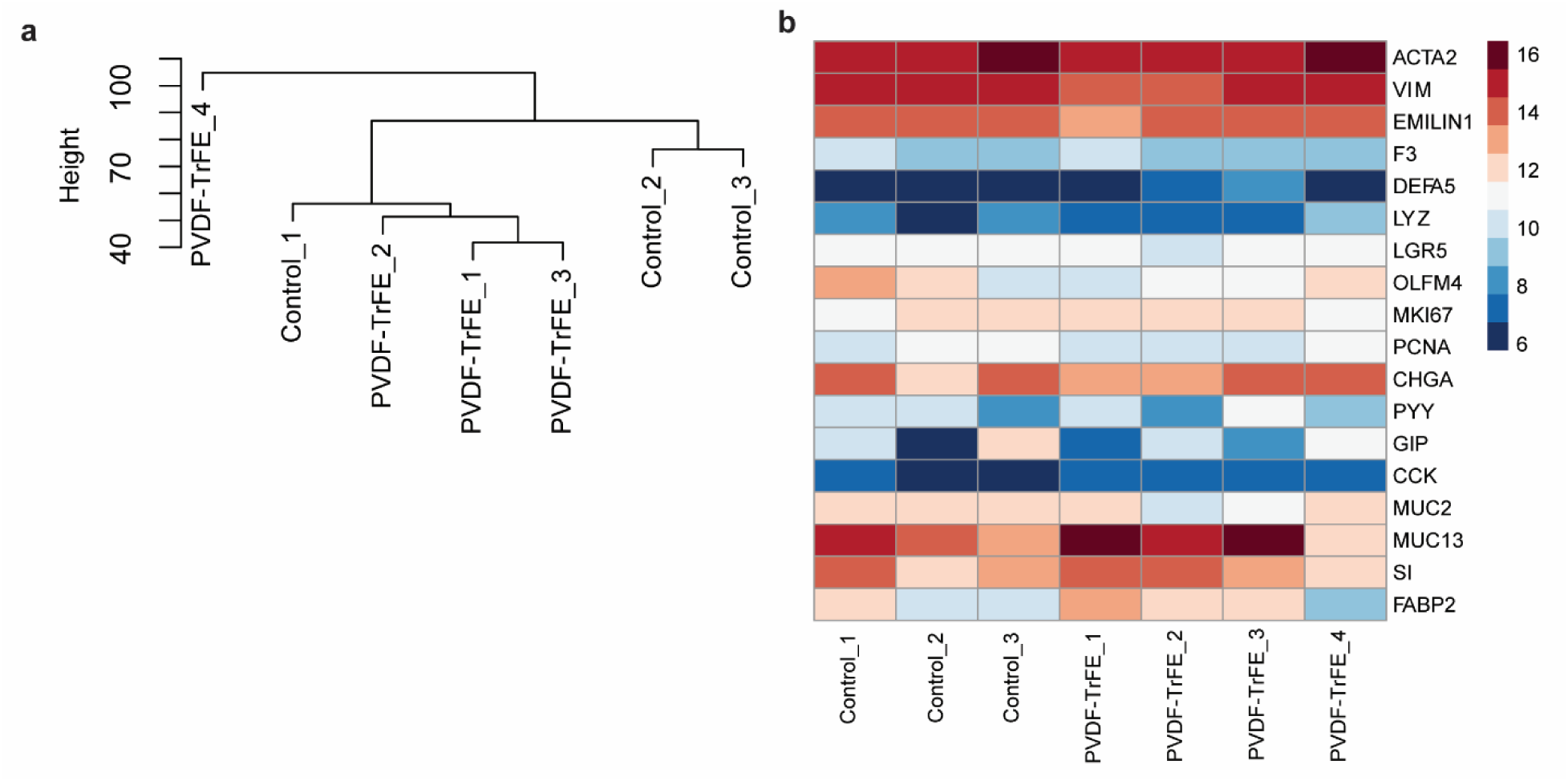
Sample dendrogram and heatmap of intestine related genes. **a**, Dendrogram of individual bulk RNA sequencing samples. **b**, Heatmap of gene expression related to intestinal cell types.

**Supplementary Figure 4.**
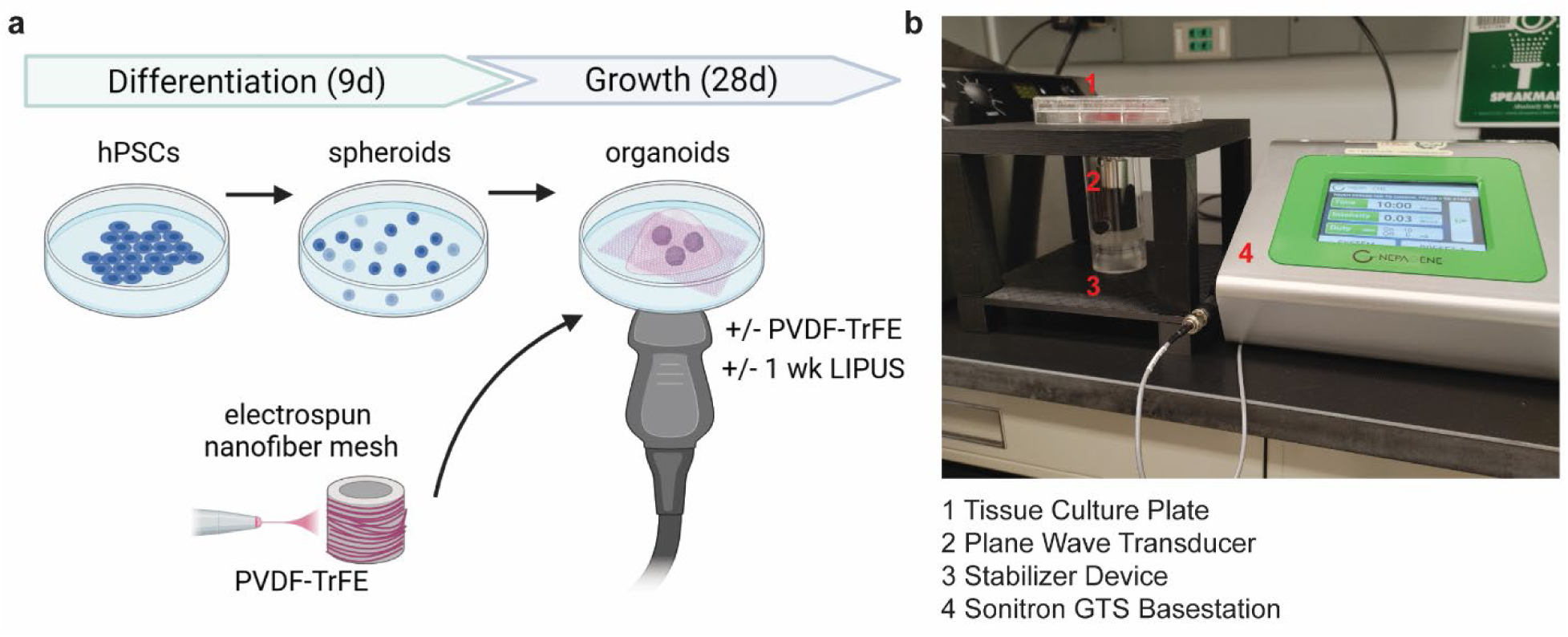
Experimental schematic for in vitro LIPUS experiment. **a**, Human pluripotent stem cells were seeded into well plates following conventional standard differentiation. After spheroids were generated and collected, they were plated with and without PVDF-TrFE as a scaffold and continued to mature for 28 days in culture. During the last week of culture, LIPUS treatments were administered every other day to HIOs with and without PVDF-TrFE, while others in both conditions remained stimulus free as scaffold condition controls. **b**, Photograph of LIPUS treatment experimental set up.

